# Effects of a Contusive Spinal Cord Injury on Spinal Motor Neuron Activity and Conduction Time in Rats

**DOI:** 10.1101/2020.06.11.146837

**Authors:** Jordan A. Borrell, Dora Krizsan-Agbas, Randolph J. Nudo, Shawn B. Frost

## Abstract

**Objective:** The purpose of this study was to determine the effects of spinal cord injury (SCI) on spike activity evoked in the hindlimb spinal cord of the rat from cortical electrical stimulation.

**Approach:** Adult, male, Sprague Dawley rats were randomly assigned to a Healthy or SCI group. SCI rats were given a 175 kDyn dorsal midline contusion injury at the level of the T8 vertebrae. At four weeks post-SCI, intracortical microstimulation (ICMS) was delivered at several sites in the hindlimb motor cortex of anesthetized rats, and evoked neural activity was recorded from corresponding sites throughout the dorsoventral depths of the spinal cord and EMG activity from hindlimb muscles.

**Main results:** In healthy rats, post-ICMS spike histograms showed reliable, evoked spike activity during a short-latency epoch 10-12 ms after the initiation of the ICMS pulse train (short). Longer latency spikes occurred between ~20-60 ms, generally following a Gaussian distribution, rising above baseline at time L_ON_, followed by a peak response (L_p_), and then falling below baseline at time L_OFF_. EMG responses occurred between L_ON_ and L_p_ (25-27 ms). In SCI rats, short-latency responses were still present, long-latency responses were disrupted or eliminated, and EMG responses were never evoked. The retention of the short-latency responses indicates that spared descending spinal fibers, most likely via the cortico-reticulospinal pathway, can still depolarize spinal cord motor neurons after a dorsal midline contusion injury.

**Significance:** This study provides novel insights into the role of alternate pathways for voluntary control of hindlimb movements after SCI that disrupts the corticospinal tract in the rat.

## Introduction

Understanding the timing of excitation of spinal cord motor neurons via descending pathways and how spinal excitability is altered by spinal cord injury (SCI) is critical for the long-range goal of functional restoration via neuromodulatory and rehabilitative interventions. While animal models of SCI have predominantly focused on cortical control of forelimb muscles, we recently established a reliable model of contusion injury in the lower thoracic spinal cord of the laboratory rat to gain insight into the capacity of spared descending fibers to participate in functional recovery of hindlimb motor behavior (Krizsan-Agbas et al., 2014; Frost et al., 2015).

The primary pathway for voluntary control of skeletal musculature in mammals is the crossed corticospinal tract (CST) originating largely from neurons in Layer 5 of the sensorimotor cortex, descending in the ventralmost part of the dorsal column (in rodents) and terminating mainly in laminae III-VI in the spinal cord dorsal horn (Casale et al., 1988; Mitchell et al., 2016). Importantly, while a few CST fibers terminate in lamina IX in the vicinity of the motor neuron cell bodies, ultrastructural studies have concluded that corticomotoneuronal connections are unsupported in rats (Yang and Lemon, 2003; Lemon, 2008). After a contusion injury to the spinal cord, cortical control of movement may be mediated by less direct polysynaptic routes from the sensorimotor cortex to spinal cord motor neurons. For example, motor cortex can influence neurons in the spinal cord indirectly via corticobulbar projections to brainstem nuclei that, in turn, project to the spinal cord. Specifically, the cells originating the reticulospinal tract (RtST), the rubrospinal tract (RbST) and the vestibulospinal tract (VST) each receive inputs from motor cortex and thus, may be substrates for post-injury motor control by the cerebral cortex. In addition, the uncrossed (ipsilateral) corticospinal tract that travels in the ventral columns of the spinal cord, is left intact by a dorsal contusion injury, and could potentially subserve motor control. The role of these spared pathways in motor control after SCI, especially with regard to the precise timing of descending supraspinal inputs, remains unclear.

Previous neurophysiological studies have evaluated corticospinal connectivity before and after SCI using transcranial magnetic stimulation (TMS) (Kamida et al., 1998; Chiba et al., 2003; Nielsen et al., 2007; Petrosyan et al., 2017), epicortical (surface) stimulation (Elger et al., 1977; Janzen et al., 1977; Ryder et al., 1991), and intracortical microstimulation (ICMS) (Bannister and Porter, 1967; Bogatyreva and Shapovalov, 1973; Stewart et al., 1990; Babalian et al., 1993; Alstermark et al., 2004). Although TMS and surface stimulation approaches are less invasive, they are spatially non-specific compared with intracortical stimulation. Due to this non-specificity in the volume of cortical tissue that is stimulated, and the multiple descending pathways giving rise to evoked responses, there are discrepancies in the published literature with regard to conduction times from cerebral cortex to spinal cord. Likewise, conduction times are often based on relatively imprecise compound action potentials (e.g., motor evoked potentials (MEPs)), that cannot discriminate peaks in activation related to depolarization of sub-populations of neurons in the spinal cord. As a result, a broad range of minimum conduction times have been reported in the literature. Reports of MEP latencies range from 1 – 14 ms in the cervical enlargement (Bannister and Porter, 1967; Stewart et al., 1990; Babalian et al., 1993; Alstermark et al., 2004; Nielsen et al., 2007) and from 2 – 35 ms in the lumbar enlargement (Bogatyreva and Shapovalov, 1973; Elger et al., 1977; Stewart et al., 1990).

Despite these limitations, examination of stimulus-evoked conduction times from cerebral cortex to spinal cord before and after SCI has provided important insights into alternative pathways for motor control after such injuries. However, the vast majority of these studies have been limited to the cervical cord. Changes in conduction times between cerebral cortex and the lumbar spinal cord have rarely been examined before and after SCI. Thus, the goal of the present study was to use the combined methodology of ICMS and intraspinal extracellular microelectrode recordings to determine the effects of a contusive SCI on the precise timing of stimulus-evoked neural activity. In the present study, we conducted neurophysiological procedures in anesthetized rats four weeks after a contusive, lower-thoracic SCI. Using a linear microelectrode array, we recorded simultaneously from multiple dorsoventral depths in the hindlimb spinal cord grey matter and clustered the recorded spikes for analysis. The resulting data allow us to describe in a more detailed, quantitative manner, the changes in the timing of ICMS-evoked spike activity after SCI. More importantly, the results provide insight into the putative descending motor pathways that may still be functional after injury. These data inform the further development of activity-dependent neuromodulation techniques that are currently being developed for human application after SCI.

## Materials and Methods

### Experimental Design

To examine the effects of a contusive SCI on cortically-evoked spinal neuronal activity, ICMS was delivered in the hindlimb area of motor cortex (HLA) and ICMS-evoked spikes and EMG signals were simultaneously recorded from hindlimb spinal cord and hindlimb muscles, respectively, in healthy and SCI rats (Figure 1).

**Figure 1.**
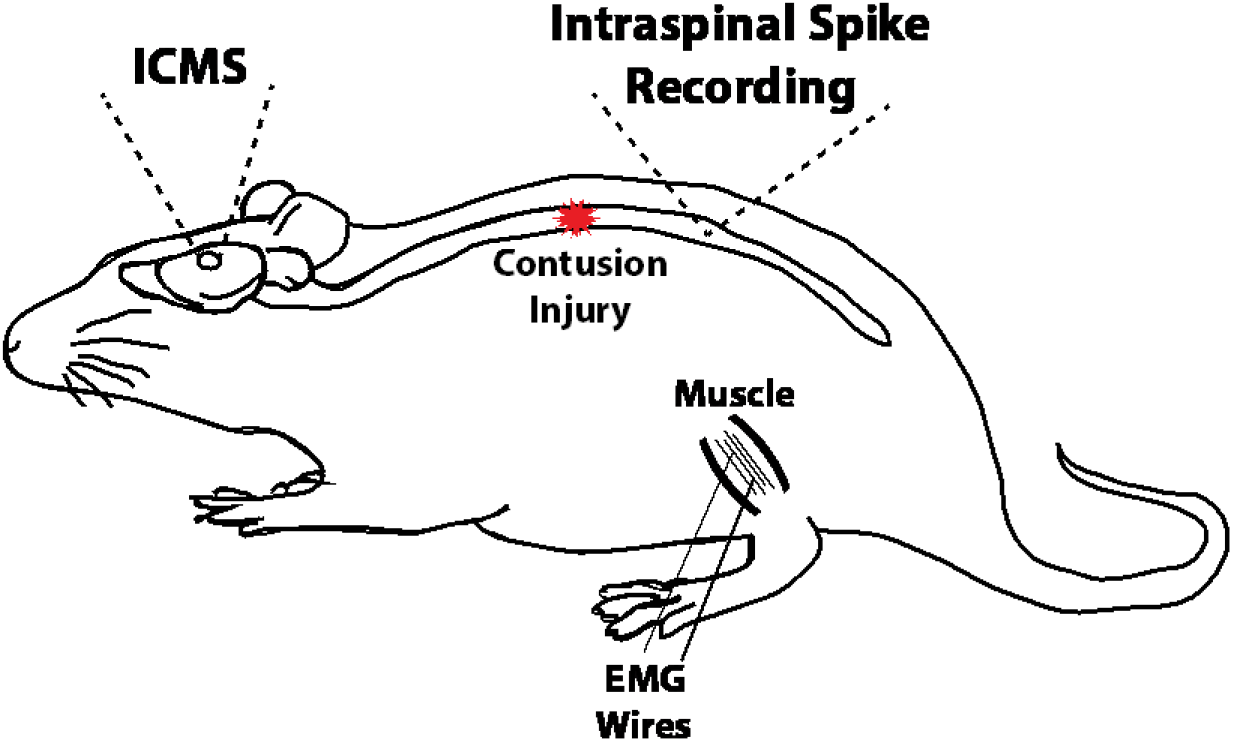
Overview of experimental design. ICMS was conducted in HLA of motor cortex in healthy and SCI rats. In SCI rats, a thoracic contusion injury was made at the level of the T8 vertebrae. Four weeks later, and under anesthesia, ICMS-evoked spikes and EMG signals were recorded simultaneously in the T13 vertebral segment (i.e., rostral lumbar enlargement) and four hindlimb muscles (EMG), respectively, for ~5 minutes during ICMS-evoked data acquisition periods.

### Subjects

A total of 20 adult, male, Sprague Dawley rats (128 ± 13.4 days old) were used in this study. Rats were randomly assigned to one of two groups: “Healthy” and “SCI”. Three rats were removed from the study due to complications during surgical procedures, resulting in a total of seven rats in the Healthy group (373.4 ± 27.27 g body weight) and ten rats in the SCI group (417.8 ± 10.24 g body weight). Rats in the SCI group underwent a surgical procedure for induction of a spinal cord contusion. All rats underwent the same electrophysiological procedures (Figure 1) described below. Average body weights at the time of the electrophysiological procedures were not statistically different between groups (*p* = 0.1043). This study was performed in accordance with all standards in the *Guide for the Care and Use of Laboratory Animals* (Institute for Laboratory Animal Research, National Research Council, Washington, DC: National Academy Press, 1996). The protocol was approved by the University of Kansas Medical Center Institutional Animal Care and Use Committee.

### Spinal Cord Contusion Procedures

Spinal cord contusion procedures followed those described in previous experiments (Krizsan-Agbas et al., 2014). SCI surgeries were performed under ketamine hydrochloride (100 mg /kg IP)/xylene (5 mg /kg IP) anesthesia in aseptic conditions. Rats in the SCI group underwent a T8 laminectomy and 175 kDyn (moderate) impact contusion injury using an Infinite Horizon spinal cord impactor (Precision Systems and Instrumentation, LLC, Fairfax Station, VA). Displacement distance reported by the impactor software for each contusion was recorded at the time of surgery and was used as an initial quantitative marker for successful impact. The average displacement value of the impactor was 1041.8 ± 111.1 μm (mean ± SD) from the surface of the spinal cord. At the conclusion of the surgery, 0.25% bupivacaine HCl was applied locally to the skin incision site. Buprenex (0.01 mg/kg, S.C.) was injected immediately after surgery and 1 day later. All animals were monitored daily until the end of the experiment. Before and after surgery, each rat received a subcutaneous injection of 30,000 U of penicillin (Combi-Pen 48). Additionally, starting the first day after surgery, daily penicillin injections were given in 5 mL saline throughout the first week to prevent infections and dehydration. Bladders were expressed twice daily until animals recovered urinary reflexes. From the second week onward, rats were supplemented with vitamin C pellets (BioServ, Frenchtown, NJ) to avert urinary tract infection.

### Behavioral Assessment Procedures

The Basso, Beattie, and Bresnahan (BBB) locomotor rating scale was used to assess locomotor ability in each of the rats (Basso et al., 1995). Initial BBB assessment was conducted in both groups to verify that rats had no pre-existing motor deficits (i.e., displayed a BBB score of 21). This initial assessment also served as a baseline measure of locomotor performance for rats in the SCI group. First, rats were habituated to the apparatus for 1-week preceding testing. In the SCI group, BBB assessment was conducted 1—3 days before SCI and at 1, 2, 3, and 4 weeks post-SCI. Briefly, rats ran along a straight alley with a darkened goal box at the end for observational purposes. Scores were recorded for the left and right sides of each animal. For SCI rats, a BBB score of 13-15 at 4 weeks post-SCI was set as the inclusion criterion for this study; however, no rats were removed from the study due to this criterion. Four weeks post-SCI was chosen for the final BBB assessment (prior to electrophysiological procedures) because BBB scores typically plateau by this SCI time point and behavior is relatively stable (Scheff et al., 2003; Krizsan-Agbas et al., 2014).

### Surgical and Electrophysiological Procedures

Electrophysiological procedures were performed 4 weeks post-SCI. Procedures took place in an RF-shielded room to prevent and reduce outside electrical interference and noise. After an initial stable anesthetic state was established using isoflurane anesthesia, the scalp and back were shaved. Then isoflurane was withdrawn, and an initial dose of ketamine hydrochloride (100 mg /kg; IP)/xylazine (5 mg/kg; IP) was administered. The anesthetic state was maintained throughout the remaining surgical and electrophysiological procedures with subsequent doses of ketamine (10 mg; IP or IM) and monitored via pinch and corneal reflexes. Additional doses of ketamine were administered if the rat reacted to a pinch of the forepaw/hindpaw, blinked after lightly touching the cornea, or exhibited increased respiration rate.

### EMG electrode implantation

EMG electrodes were implanted in four hindlimb muscles: the lateral gastrocnemius (LG), tibialis anterior (TG), vastus lateralis (VL), and biceps femoris (BF) (Borrell et al., 2017). Briefly, each EMG electrode consisted of a pair of insulated multi-stranded stainless-steel wires exposed approximately 1 mm, with the implanted end of the wire folded back on itself (‘hook’ electrode). Implantation locations were determined by surface palpation of the skin and underlying musculature. Once the hindlimbs were shaved, EMG electrodes were inserted into the belly of each muscle with the aid of a 22-gauge hypodermic needle. For each EMG electrode pair, wires were positioned approximately 5 mm apart in each muscle. The external portion of the wires was secured to the skin with surgical glue (3M Vetbond Tissue Adhesive, St. Paul, MN) and adhesive tape. An additional ground lead was placed into the base of the tail.

To verify that the EMG electrodes were in the belly of the target muscle, a stimulus isolator (BAK Electronics, Inc., Umatilla, FL) was used to deliver a biphasic (cathodic-leading) square-wave current pulse to the muscle through the implanted EMG electrodes. In addition, the impedance between the EMG electrodes was tested via an electrode impedance tester (BAK Electronics, Inc., Umatilla, FL). The EMG electrodes were determined to be inserted properly and in the desired muscle if: (a) the electrode impedance was approximately 7-8 kΩ; (b) direct current delivery to the muscle resulted in contraction of the desired muscle, and (c) the movement threshold was ≤ 5mA.

### Intracortical microstimulation

The rats were placed in a Kopf small-animal stereotaxic frame (David Kopf Instruments®, Tujunga, CA) and the incisor bar was adjusted until the heights of lambda and bregma were equal (flat skull position). The cisterna magna was punctured at the base of the skull to reduce edema during electrophysiological procedures. A craniectomy was performed over the hindlimb motor cortex (HLA). The general location of the craniectomy was guided by previous motor mapping studies in the rat (Frost et al., 2013; Frost et al., 2015). The dura over the cranial opening was incised and the opening was filled with warm, medical grade, sterile, silicone oil (50% Medical Silicone Fluid 12,500, 50% MDM Silicone Fluid 1000, Applied Silicone Corp., Santa Paula, CA) to prevent desiccation during the experiment.

A magnified digital color photograph of the exposed cortex was taken through a surgical microscope (Figure 2A and 2B). The image was transferred to a graphics program (Canvas 3.5) where a 200 μm grid was superimposed, by calibration with a millimeter ruler, to indicate intended sites for microelectrode penetration. The 0.0 mm rostrocaudal and mediolateral reference point was defined by the location of bregma.

**Figure 2.**
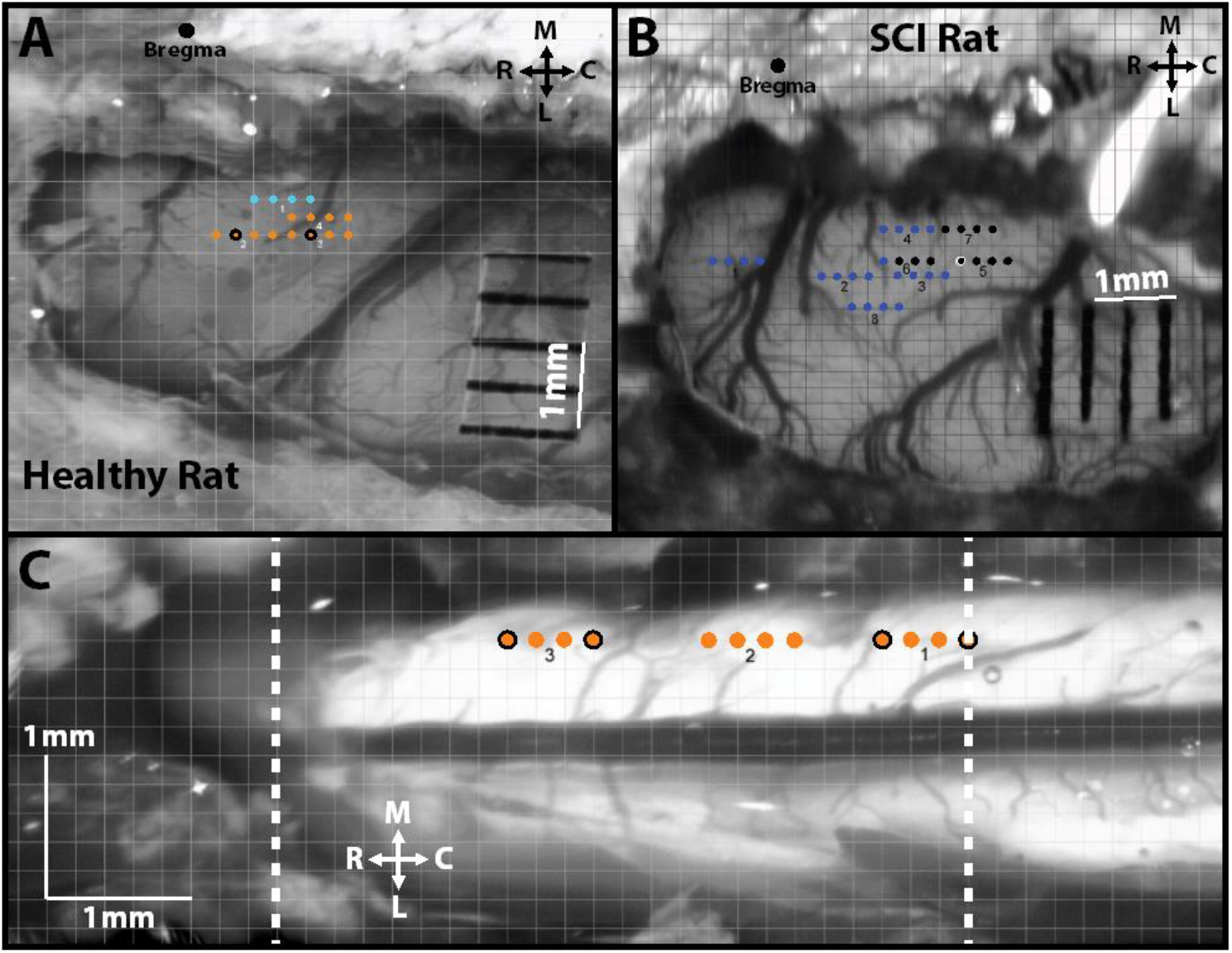
Overview of ICMS and intraspinal microstimulation (ISMS) mapping procedures. **A)** Cranial opening in a healthy rat over HLA with superimposed grid for ICMS mapping. The solid black dot represents Bregma; each orange dot represents a site where ICMS evoked a hindlimb movement. Each light blue dot represents a site where ICMS evoked a trunk movement. **B)** Cranial opening in a SCI rat over HLA with superimposed grid for ICMS mapping. The larger solid black dot represents Bregma; each small solid black dot represents a site where ICMS did not evoke a movement. Each blue dot represents a site where ICMS evoked a forelimb movement. **C)** Spinal cord opening in a healthy rat over the hindlimb spinal cord with superimposed grid for ISMS mapping. The two dashed white lines indicate the rostral-caudal extent of the T13 vertebra. Each orange dot represents a site where ISMS evoked a hindlimb movement. **A, B, C)** Orange dots with black outline and black dot with white outline indicate sites with similarly evoked movements that were paired for analysis. Each square box is 200 x 200 μm. R = Rostral, C = Caudal, M = Medial, and L = Lateral.

ICMS was conducted in HLA using an activated single-shank, 8-site Neuronexus probe (Neuronexus, Ann Arbor, MI) optimized for microstimulation, following standard procedures (Frost et al., 2013; Frost et al., 2015). ICMS was delivered at regularly spaced sites on the superimposed 200 μm grid at a depth of ~1.7 mm below the cortical surface (i.e., in Layer 5) from the ventral-most site on the probe. Electrode depth was controlled using a Kopf hydraulic microdrive (Kopf Instruments, Tujunga, CA). To determine ICMS-evoked movements, stimuli consisted of thirteen, 200 μs biphasic cathodal pulses delivered at 300 Hz repeated at 1/sec from an electrically isolated, charge-balanced (capacitively coupled) stimulation circuit. Each activated stimulation site had an impedance in the range of ~40 – 60 kΩ with a surface area of 1250 μm^2^. The maximum compliance voltage of the system was 24 V.

#### ICMS-evoked movements

At each cortical site, skeletal movement that was evoked at the threshold ICMS current intensity (i.e., movement threshold) was recorded for each stimulation site. The threshold current intensity was defined as the intensity at which a movement was evoked in ≥50% of the pulse trains. If the movement was restricted to a specific joint, it was then categorized as a hip, knee, ankle, or digit movement with a further categorization of flexion, extension, inversion, eversion, abduction, or adduction motion of the joint. ICMS current intensity did not exceed 100 μA to minimize current spread. If a movement was not evoked at a particular site, the site was labeled as non-responsive.

#### ICMS-evoked EMG

Using custom software (Matlab; The Mathworks, Inc., Natick, MA), stimulus-triggered averages (StTAs) of rectified EMG of were analyzed to determine muscle activation (Hudson et al., 2015; Borrell et al., 2017). For each of the implanted muscles, EMG signals were recorded for at least 10 ICMS stimulus trains to obtain StTAs over a 220 ms time window. In healthy rats, the current intensity used for obtaining StTAs was the threshold current intensity. Since ICMS-evoked hindlimb movements could not be evoked in SCI rats, even at 100 μA, the current intensity for ICMS-evoked EMG was set at the average movement threshold seen in the healthy rats, but not > 80 μA. Using StTAs, EMG recordings provided a more sensitive measure compared with evoked movements for evaluating motor neuron activation of skeletal muscles. StTAs were aligned to the time of the first stimulus pulse (i.e., 0 ms) and included data from −20.2 to +199.9 ms relative to the time of the first stimulus. The first 20 ms of the StTA (−20.2 to 0 ms) was defined as the baseline EMG period. A muscle was considered to be “active” when the average EMG during the post-ICMS period (0 ms to 199.9 ms) reached a peak ≥ 2.25 SD above the average EMG during the baseline EMG period and remained above this level for ≥ 3 ms. The stimulus artifact was minimal to absent in EMG recordings with no muscle activation. EMG potentials were high- and low-pass filtered (30 Hz-2.5 kHz), amplified 200-1,000 fold, digitized at 5 kHz, rectified and recorded on an RX-8 multi-channel processor (Tucker-Davis Technology, Alachua, FL).

### Intraspinal microstimulation mapping

Following ICMS mapping, a midline incision was made over the spinal column, and a laminectomy was performed on the T13-L1 vertebrae exposing the L2-S1 segments of the spinal cord. Previously derived 3-dimensional ISMS-evoked topographic maps were used to guide the location of the spinal opening to obtain stimulation sites in the hindlimb spinal cord (Padmanabhan and Singh, 1979; Borrell et al., 2017). The dura mater was removed using fine forceps and small scissors to allow microelectrode penetration. The spinal column was stabilized with a custom rodent spinal fixation frame (Keck Center for Collaborative Neuroscience Rutgers, The State University of New Jersey, USA) attached to the dorsal processes of vertebrae T12 and L2.

For intraspinal microstimulation (ISMS) maps a digital photograph of the hindlimb spinal cord (i.e., T13-L1 vertebrae) was transferred to a graphics program (Canvas 3.5) where a 200 μm grid was superimposed to indicate intended sites for microelectrode penetration (Figure 2C). Stimulation sites were ~0.8 mm lateral of the central blood vessel (i.e., midline) at a depth ~2.27 mm below the surface of the spinal cord (i.e., in the ventral horn). Each stimulus consisted of three, 200 μs biphasic cathodal pulses delivered at 300 Hz repeated at 1/sec. These ISMS parameters are commonly used for ISMS mapping procedures (Sunshine et al., 2013; Shahdoost et al., 2014a; Borrell et al., 2017).

#### ISMS-evoked movements

Movements that were evoked at the lowest intensity of ISMS stimulation (i.e., movement threshold) were recorded for each stimulation site as described previously. Sites with comparable movements that were evoked by in ICMS in HLA and ISMS in the ventral horn of the hindlimb spinal cord, respectively, were selected for neurophysiological assessment (e.g., ICMS and ISMS sites both resulting in hip flexion were compared).

#### ISMS-evoked EMG

Recording and processing of EMG data paralleled those described above for ICMS-evoked EMG, except that the ISMS train consisted of only three pulses, further minimizing the effects of the stimulus artifact on the recorded data.

### Extracellular spike recording in spinal cord

#### Microelectrode probe placement

Microelectrode placement in the cortex (stimulating electrode) and spinal cord (recording electrode) was based on the topography of the ICMS- and ISMS-movement maps (Figure 2). Microelectrodes were placed in sites where movements about the hip were evoked in both the cortex and spinal cord (orange dots in Figures 2A and 2C). Since hindlimb responses cannot be evoked from HLA in SCI rats (Frost et al., 2015), the microelectrode was placed in a non-responsive site (black dot in Figure 2B) that was directly caudal to the boundary of the forelimb motor cortex (blue dots in Figure 2B). The same stimulating microelectrode used during ICMS mapping was placed in the HLA and a recording microelectrode probe simultaneously placed in the hindlimb movement-matched spinal cord site. The recording microelectrode was a single-shank, 16-channel Michigan-style linear microelectrode probe (Neuronexus, Ann Arbor, MI), optimized for extracellular spike recording (Figure 3). Each of the 16 electrode sites had a site area of 703 μm^2^, separation of 100μm, shank thickness of 15 μm in diameter, and impedance of ~0.3 MΩ. The tip of the probe was lowered to a depth of ~2270 μm below the spinal cord surface. For analysis of spike responses as a function of depth, the 16 recording channels were divided into four sectors based on depth below the surface of the spinal cord: Dorsal Sector (720—1020 mm), Upper Intermediate Sector (1120—1420 mm), Lower Intermediate Sector (1520—1820 mm), and Ventral Sector (1920—2220 mm). With respect to spinal cord grey laminar organization, these sectors correspond to the dorsal horn, the upper part of the intermediate grey, the lower part of the intermediate grey, and the ventral horn.

**Figure 3.**
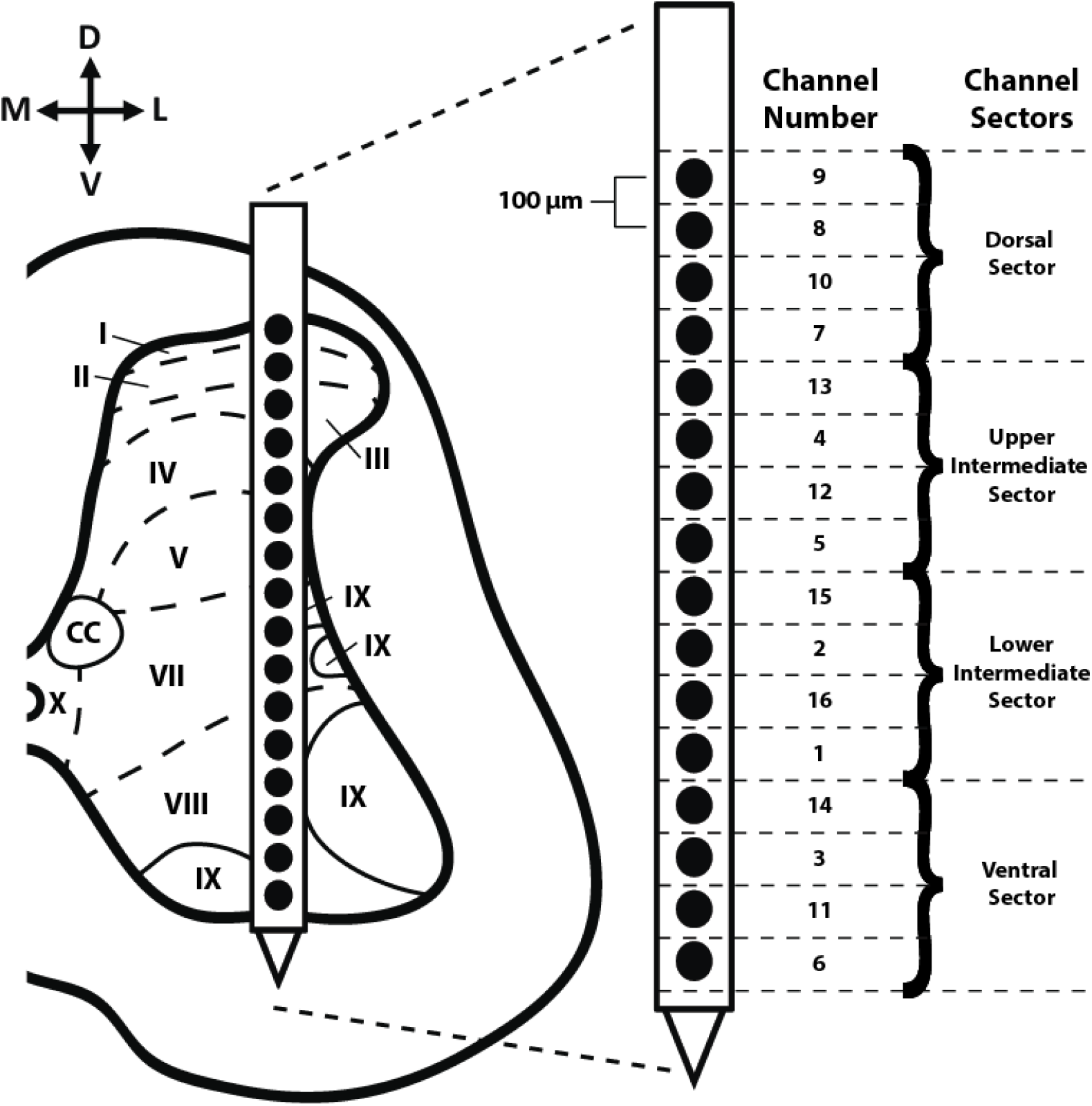
Cross sectional diagram of hindlimb spinal cord with placement of the recording probe during ICMS. ICMS-evoked extracellular spikes were recorded with a 16-channel Neuronexus recording probe inserted into the hindlimb spinal cord under T13 vertebrae ~0.8 mm from midline to a depth of ~2.27 mm below the surface of the spinal cord spanning virtually the entire dorsoventral extent of the spinal cord grey matter from the dorsal horn to the ventral horn. The schematic diagram of the recording shank indicates the channel numbers of the recording sites and corresponding dorsoventral sectors. Approximate anatomical locations of Rexed laminae are shown relative to each recording channel.

The ability of spinal cord neurons to be depolarized by cortical stimulation, hereafter called “corticospinal coupling”, was assessed using a train of three, 200 μs biphasic cathodal pulses delivered at 300 Hz to the HLA, while simultaneously recording extracellular spike events from the 16 microelectrode channels in the spinal cord and EMG from the hindlimb muscles. The stimulus train was repeated at the rate of 1 Hz, and cumulative post-stimulus spike histograms were derived. Restricting ICMS to three stimulation pulses prevented the influence of movement artifacts in the recorded data. Also, the minimal pulse train minimized the effects of stimulus artifacts on our ability to discriminate spikes. A maximum current of 100 μA was used for sites in which no movement was evoked. Corticospinal coupling sessions were conducted for 5 min at each ICMS test site.

#### Spike recording and analysis

ICMS-evoked spike activity was recorded from each of the 16 active recording channels for ~5 min for each ICMS site using neurophysiological recording and analysis equipment (Tucker Davis Technologies, Alachua, FL). Neural spikes were discriminated using principle component analysis (Lewicki, 1998). ICMS-evoked spike activity was sorted and averaged into 1 ms bins over the ~5 min recording bout (i.e., averaged over ~300 ICMS trains). Activity was represented in post-stimulus spike histograms for 200 ms around the onset of each ICMS stimulus (ICMS began at 0 ms; histogram extended −100 ms before and +100 ms after onset ICMS), using custom software (Matlab; The Mathworks, Inc., Natick, MA) (Riddle and Baker, 2010). Spike activity could not be discriminated over the first ~7 ms due to the stimulus artifact from the three stimulus pulses.

#### Firing rate (FR) calculation

To quantitatively analyze these data and compare Healthy and SCI groups, the ICMS-evoked firing rate (FR) (i.e., total number of recorded spikes per 1ms bin/total time of recording) was calculated within each time bin of the post-stimulus spike histograms using custom software (Matlab; The Mathworks, Inc., Natick, MA). The FR was only considered for analysis if the FR for the respective time bin was greater than 2x standard deviation (horizontal, dashed, red line; Figure 6) above average baseline (horizontal, solid, red line; Figure 6) FR. Baseline FR was derived by averaging the FR of each time bin 10 ms before the onset of ICMS (i.e., from −10 to 0 ms in the post-stimulus spike histogram).

#### Probability of activation

The Probability of Activation (POA) is defined as the likelihood that spikes are evoked during each ICMS stimulus event. The POA of ICMS-evoked spikes occurring during each test stimulus of ICMS was calculated for each 5-min recording. POA = number of recorded ICMS-evoked spiking incidences/total number of ICMS-test stimuli. The POA was calculated for each recording channel at each specified latency and compared between Healthy and SCI groups and among dorsoventral sectors. The mean and standard error were calculated by averaging the POA of each channel within each respective dorsoventral sector.

### Euthanasia & Histology

At the end of each experiment, animals were euthanized with an intraperitoneal injection of sodium pentobarbital (Beuthanasia-D; 100 mg kg^−1^) and transcardially perfused with 4% paraformaldehyde in 0.1 M PBS. The spinal cord was sectioned at 30 μm on a cryostat in the coronal plane. Histology was performed on selected sections with cresyl violet stain for Nissl bodies to verify the extent of the injury.

### Statistical Analyses

Statistical analyses were performed using JMP 11 software (SAS Institute Inc., Cary, NC). Since the temporal distribution of spike firing patterns tends to have a Poisson rather than Gaussian distribution, the data were evaluated using a nonparametric Mann-Whitney U test to compare the medians of each dorsoventral sector between Healthy and SCI rats. Other endpoints that were normally distributed were evaluated using a student’s t-test to compare means of each dorsoventral sector between healthy and SCI rats and are presented as average ± standard error of the mean unless otherwise stated.

## Results

### BBB Scores

Each of the rats in the Healthy group scored 21 on the BBB scale (normal) while the rats in the SCI group had a significantly lower average BBB score of 14.7 ± 0.8 (mean ± SD) at 4 weeks post-SCI (*p* < 0.0001). This lower BBB score indicates that the SCI rats had consistent weight-supported plantar steps, no or occasional toe clearance (i.e., scraping of the digits on the ground) during forward limb advancement, consistent forelimb-hindlimb coordination, and predominant paw position during locomotion that was always rotated externally just before it lifted off at the end of stance and either externally rotated or parallel to the body at initial contact with the surface.

### SCI Verification

An exemplar coronal histological section through the center of the injury is shown in Figure 4A. After SCI, the spinal grey matter at the epicenter was severely damaged. The ventromedial (location of the ventral CST and RtST) and dorsolateral (location of the RbST and dorsolateral CST) spinal cord white matter tracts generally remained intact while the dorsal column white matter tracts (location of the dorsomedial CST in rodents) was severely damaged (Figure 4B).

**Figure 4.**
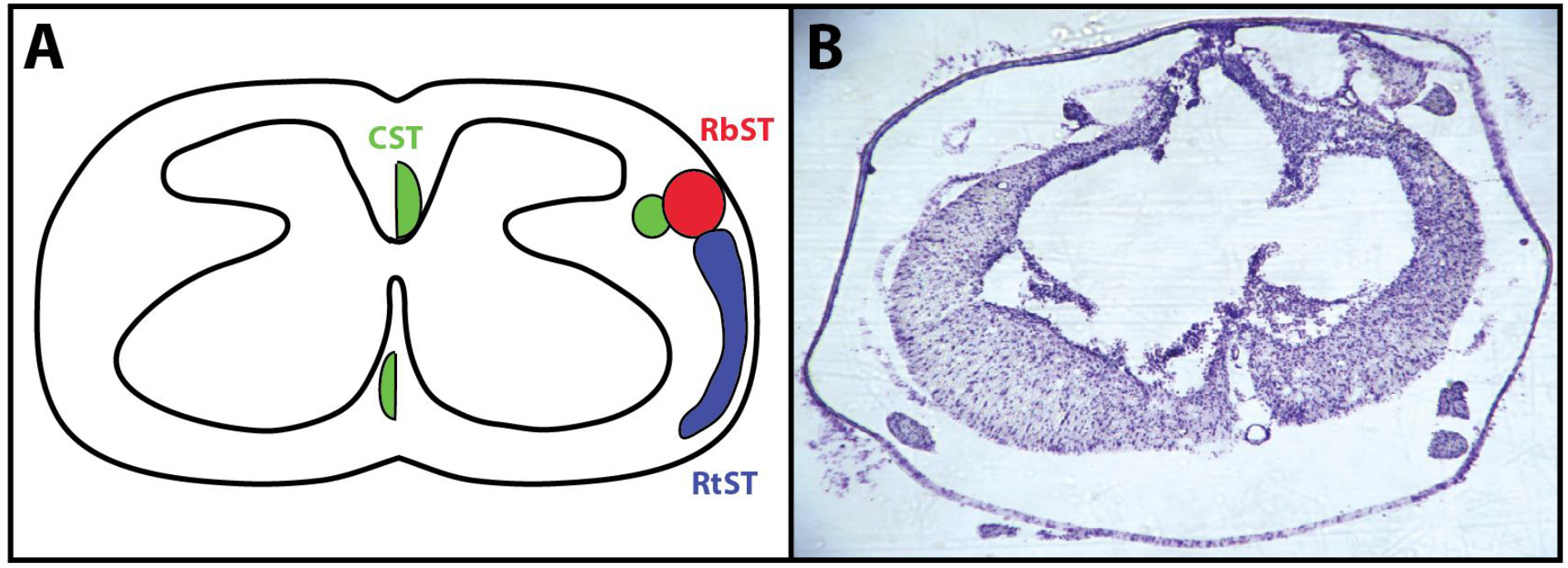
Verification of spinal cord injury under T8 vertebrae. A) Schematic diagram of a coronal section identifying the location of the descending corticospinal tract (CST; green), rubrospinal tract (RbST; red), and reticulospinal tract (RtST; blue) (Lemon, 2008; Fink and Cafferty, 2016). B) Representative image of spinal cord injury epicenter stained with cresyl violet 4 weeks after injury.

### Neurophysiological Results

Neurophysiological sessions were conducted in seven rats in the Healthy group (56 cumulative sessions) and ten rats in the SCI group (10 cumulative sessions; 4 weeks post-SCI). In Healthy rats, ICMS-evoked activity was examined at multiple sites in separate data collection sessions. Due to the inability to evoke hindlimb movements via ICMS after SCI, only one session at one cortical site was conducted per SCI rat. The cortical stimulation site selected in SCI rats was based on the location of ICMS-evoked hip movement sites from ICMS maps in healthy rats. In Healthy rats, hip movement sites are consistently located caudal to the forelimb representation and caudolateral to the trunk representation (Frost et al., 2015). Both forelimb and trunk movements can still be evoked after lower thoracic SCI. The average amount of ketamine (mean ± SD) administered during the experimental surgery was not significantly different between groups (Healthy = 122.14 ± 17.66 mg; SCI = 119.30 ± 24.03 mg; *p* = 0.7941).

### ICMS-evoked movements and EMG onset latencies

In Healthy rats, evoked movements from ICMS in HLA were observed as described in previous studies. Evoked responses consisted of hip flexion (n = 23 sites), lateral leg rotation (n = 5 sites), knee flexion (n = 8 sites), ankle flexion (n = 9 sites), digit extension (n = 6 sites), and no response (n = 5 sites). In SCI rats, no ICMS-evoked hindlimb movements were elicited. The average ICMS movement threshold current intensity was 80.66 ± 31.78 μA (mean ± SD) in Healthy rats. Based on the average ICMS movement threshold seen in Healthy rats, a constant ICMS current amplitude of 80 μA was used in SCI rats. No significant differences in ICMS threshold currents were found among the different movement categories (*p* > 0.05). Thus, data from the different hindlimb movement categories were pooled in subsequent analyses of ICMS-evoked activity in the spinal cord.

ICMS-evoked EMG potentials were recorded from four hindlimb muscles: Biceps Femoris (BF) Vastus Lateralis (VA), Tibialis Anterior (TA), and Lateral Gastrocnemius (LG). Onset latencies were determined from stimulus-triggered averages (StTAs) of each ICMS recording (Figure 5 Left). ICMS-evoked EMG onset latencies (i.e., 2.25 standard deviations above baseline) from Healthy rats (Figure 5 Left) were as follows (mean ± standard error): BF = 26.30 ± 1.00 ms, VL = 26.27 ± 1.23 ms, TA = 27.21 ± 1.03 ms, and LG = 25.82 ± 0.94 ms. Post hoc comparisons (student’s t-test) showed no significant differences in mean EMG latencies when comparing ICMS-evoked muscles. After SCI, ICMS-evoked EMG activity could not be elicited (Figure 5 Right).

**Figure 5.**
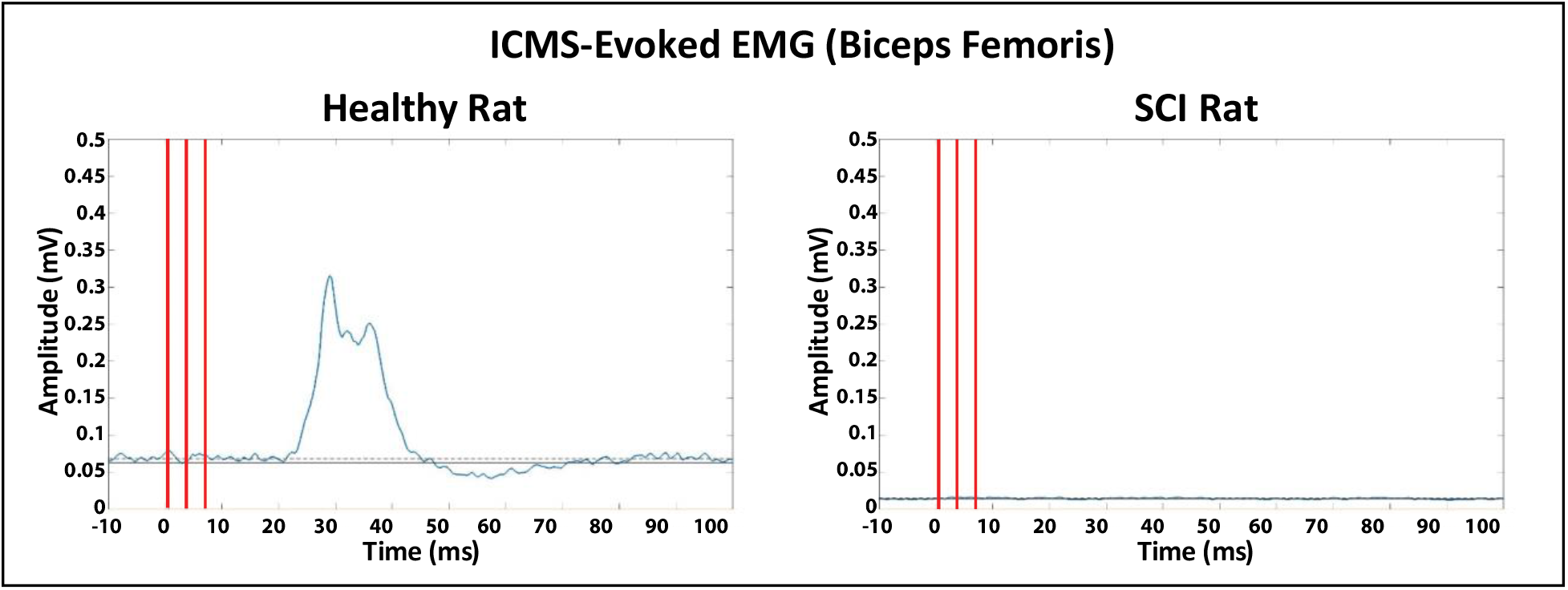
Representative ICMS-evoked EMG recording from the Biceps Femoris of a Healthy rat (left) and an SCI rat (right). Stimulus-triggered averaging (StTA) was used to analyze the EMG recordings. Latency was measured as the first instance a muscle was active after onset stimulation at 2.25 standard deviations above baseline average (horizontal dashed line). Vertical red lines are the superimposed stimulus artifacts from the 3-pulse stimulation train. Ten stimulus trains were averaged for StTA analysis.

### ICMS-evoked spike activity in spinal cord neurons

#### Overview of spike latencies

ICMS-evoked spikes in the spinal cord (Figure 6A) were recorded extracellularly, sorted, and displayed in post-stimulus time histograms. Healthy rats typically displayed two populations, differing in onset latency. Short latency spikes typically occurred between 10-12 ms post-ICMS (Figure 6B). Long latency spikes were distributed over a broader Gaussian distribution (~20-60 ms) after the short latency responses. Long latency spikes were divided into “long-latency onset” (L_on_; i.e., onset of Gaussian distribution; rising above 2x standard deviation of average baseline activity), “peak latency” (L_p_; i.e., peak of Gaussian distribution), and “long-latency offset” (L_off_; i.e., offset of Gaussian distribution; falling below 2x standard deviation of average baseline activity) (Figure 6B). L_off_ latencies were not consistent and displayed high variability (mean ± standard deviation; Dorsal Sector = 61.73 ± 26.90 ms, Upper Intermediate Sector = 66.86 ± 27.17 ms, Lower Intermediate Sector = 63.87 ± 23.89 ms, and Ventral Sector = 63.95 ± 24.01 ms). Thus, L_off_ latencies were not considered for further analysis.

**Figure 6.**
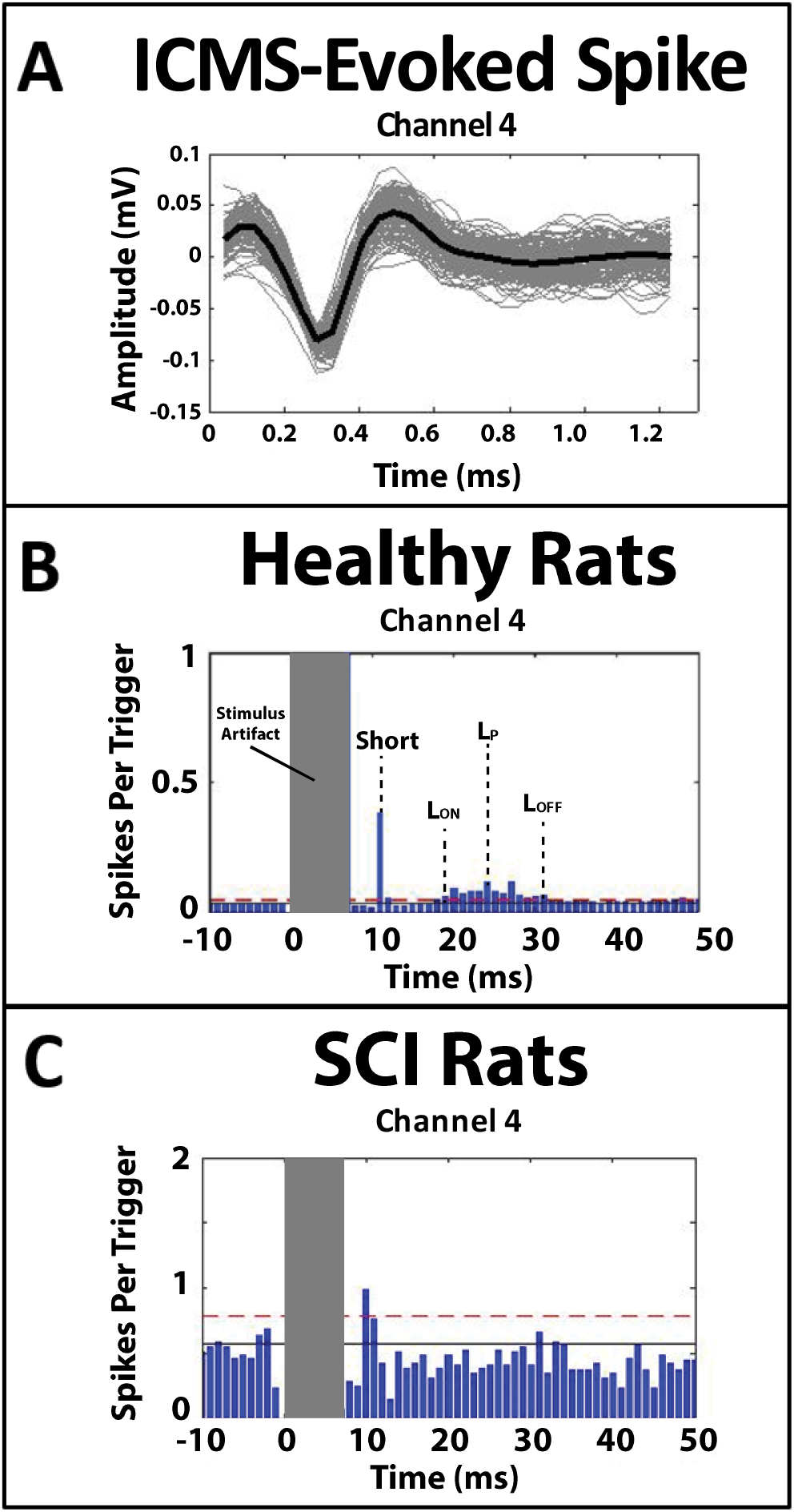
ICMS-evoked activity in the spinal cord. Sample ICMS-evoked action potentials (i.e., spikes) recorded from hindlimb spinal cord and post-stimulus spike histogram analysis from the summation of all rats in recording channel 4 (i.e., depth of 1220 μm). Data were normalized to the largest spiking bin. **A)** ICMS-evoked spikes recorded from one channel in an intermediate layer of hindlimb spinal cord. Solid black spike is the average of all recorded spikes from channel 4, and grey spikes are all spikes of similar spiking profiles recorded from channel 4. **B)** Post-stimulus spike histogram of spikes recorded from channel 4 from all healthy rats. Solid horizontal line is baseline average. Red dashed horizontal line is 2 standard deviations (SDs) above baseline average. Onset of ICMS stimulation is at 0 ms. Spinal cord spiking activity was seen at two different latencies, which were separated into short- and long-latency times. “Short” is the short-latency time at ~10 ms. L_on_ is onset time when long latency evoked spiking activity reaches above 2 SDs above baseline average, L_p_ is the peak of long latency evoked spiking activity, and L_off_ is offset time before long latency spiking activity drops below 2 SDs above baseline average. The stimulus artifact occurred from 0-7 ms. **C)** Post-stimulus spike histogram of spikes recorded from channel 4 of all SCI rats. Short latency spikes consistently occurred in all dorsoventral sectors. The solid horizontal line is the baseline average. Red dashed horizontal line is 2 standard deviations (SDs) above baseline average. The long latency responses were greatly diminished in all dorsoventral sectors after SCI.

#### Post-ICMS spike histograms at different dorsoventral depths

For each rat in both Healthy and SCI groups, ICMS-evoked spikes were sorted and examined in post-stimulus spike histograms organized by channel (dorsoventral depth). Not all recorded latencies (Short, L_on_, L_p_, and L_off_) were observed in all channels (Figure 7), nor in each rat; particularly in the SCI rats, nor was this absence of observed latencies specific to a particular dorsoventral depth. In the seven healthy rats, evoked spikes occurred at the Short latency in six rats (86%). In nine of the 10 SCI rats, evoked spikes occurred at the Short latency (90%). Representative recordings from a single corticospinal coupling session between cortex and spinal cord in one Healthy and one SCI rat are shown in Figure 7. In the Healthy rat in this example, two channels did not display any long-latency responses. The SCI rat in this example showed diminished evoked activity compared with the healthy rat; however, in nine channels ICMS-evoked activity occurred at the Short latency.

**Figure 7.**
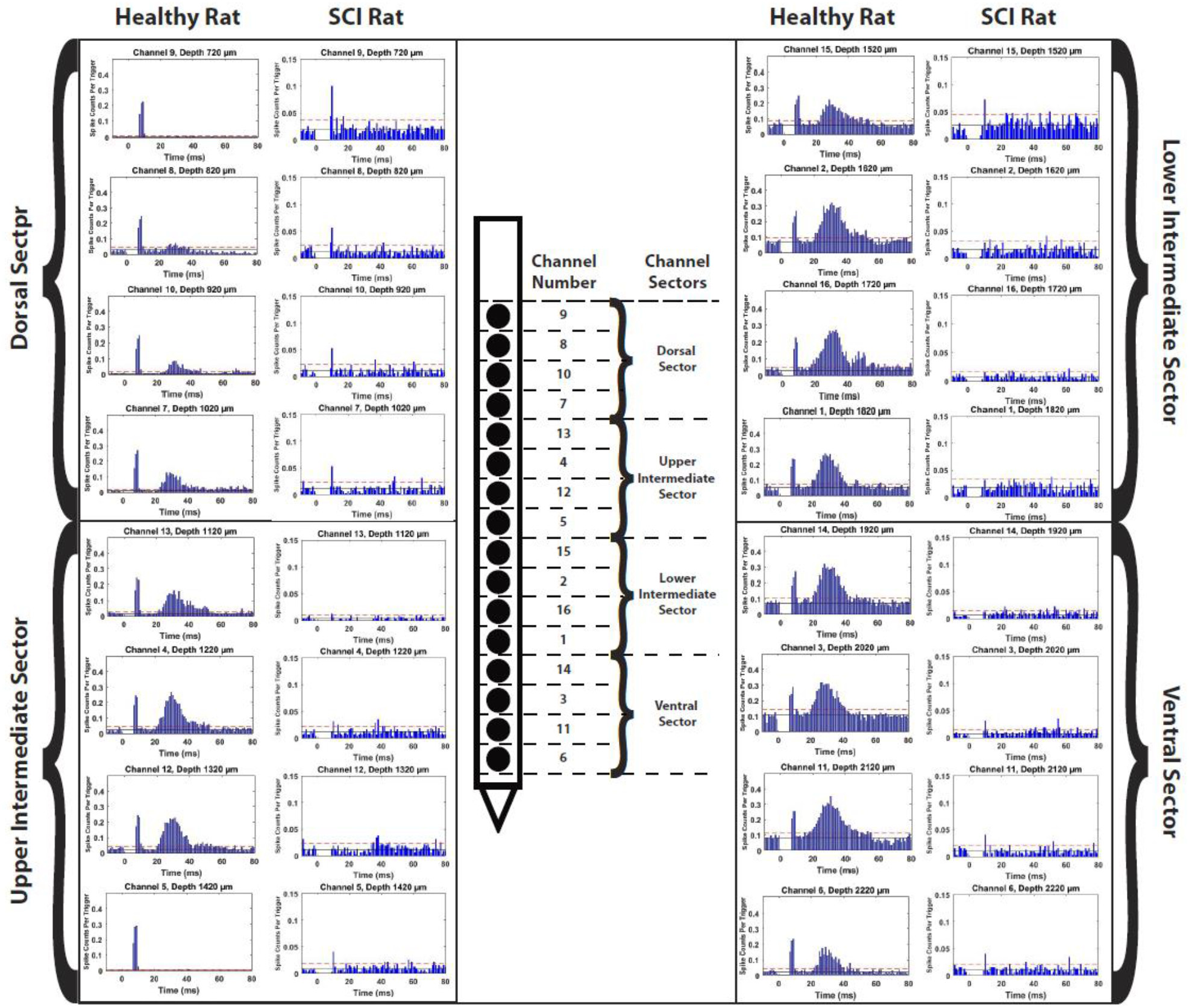
Qualitative comparison of ICMS-evoked activity. Representative recordings from one Healthy and one SCI rat from multiple recording channels (i.e., dorsoventral depths) divided into dorsoventral sectors based on depth. Qualitatively, the long latency responses are diminished in the SCI rat while the short latency response appears to remain present in the SCI rat compared to the Healthy rat.

#### Probability of activation (POA)

The POA for Healthy and SCI rats was calculated for each dorsoventral sector and listed in Table 1. At the Short latency, the POA significantly increased in all dorsoventral sectors after SCI (between-group comparisons; Dorsal Sector, *p* = 0.0001; Upper Intermediate Sector, *p* = 0.0004; Lower Intermediate Sector, *p* = 0.0081; Ventral Sector, *p* = 0.0004). At the L_on_ latency, the POA significantly increased in the Lower Intermediate (*p* = 0.0138) and Ventral (*p* = 0.0001) Sectors after SCI. At the L_p_ latency, the POA significantly decreased in the Upper Intermediate (*p* = 0.0187) Sector and significantly increased in the Ventral (*p* < 0.0001) Sector after SCI.

**Table 1.** The Probability of Activation (POA) of ICMS-evoked spikes at each specified latency. The POA was calculated for each group for each dorsoventral sector. A green up-arrow indicates a significant increase in POA after SCI while a red down-arrow indicates a significant decrease in POA after SCI. **\***= Significant difference (*p* < 0.05) when compared to healthy rats of the same dorsoventral sector and specified latency.

| Dorsoventral Sector | Rat Group | Probability of Activation (Mean ± SE) | | | | | |
| --- | --- | --- | --- | --- | --- | --- | --- |
| | | Short | | $L_{on}$ | | $L_p$ | |
| Dorsal | Healthy | 0.22 ± 0.03 | ↑ | 0.72 ± 0.03 | = | 0.78 ± 0.03 | = |
| | SCI | 0.48 ± 0.03* | | 0.73 ± 0.05 | | 0.70 ± 0.04 | |
| Upper Intermediate | Healthy | 0.27 ± 0.02 | ↑ | 0.90 ± 0.02 | = | 0.93 ± 0.02 | ↓ |
| | SCI | 0.50 ± 0.04* | | 0.85 ± 0.05 | | 0.78 ± 0.09* | |
| Lower Intermediate | Healthy | 0.18 ± 0.02 | ↑ | 0.64 ± 0.09 | ↑ | 0.66 ± 0.08 | = |
| | SCI | 0.35 ± 0.09* | | 0.80 ± 0.04* | | 0.75 ± 0.03 | |
| Ventral | Healthy | 0.19 ± 0.02 | ↑ | 0.54 ± 0.07 | ↑ | 0.56 ± 0.05 | ↑ |
| | SCI | 0.43 ± 0.03* | | 0.80 ± 0.06* | | 0.83 ± 0.03* | |

#### Baseline firing rate

The median baseline firing rate (before onset of ICMS) was calculated for Healthy and SCI rats and is displayed in Figure 8. The median baseline firing rate was significantly higher in the Upper Intermediate Sector (*p* = 0.0433) and significantly lower in the Ventral (*p* = 0.0.0050) Sector after SCI.

**Figure 8.**
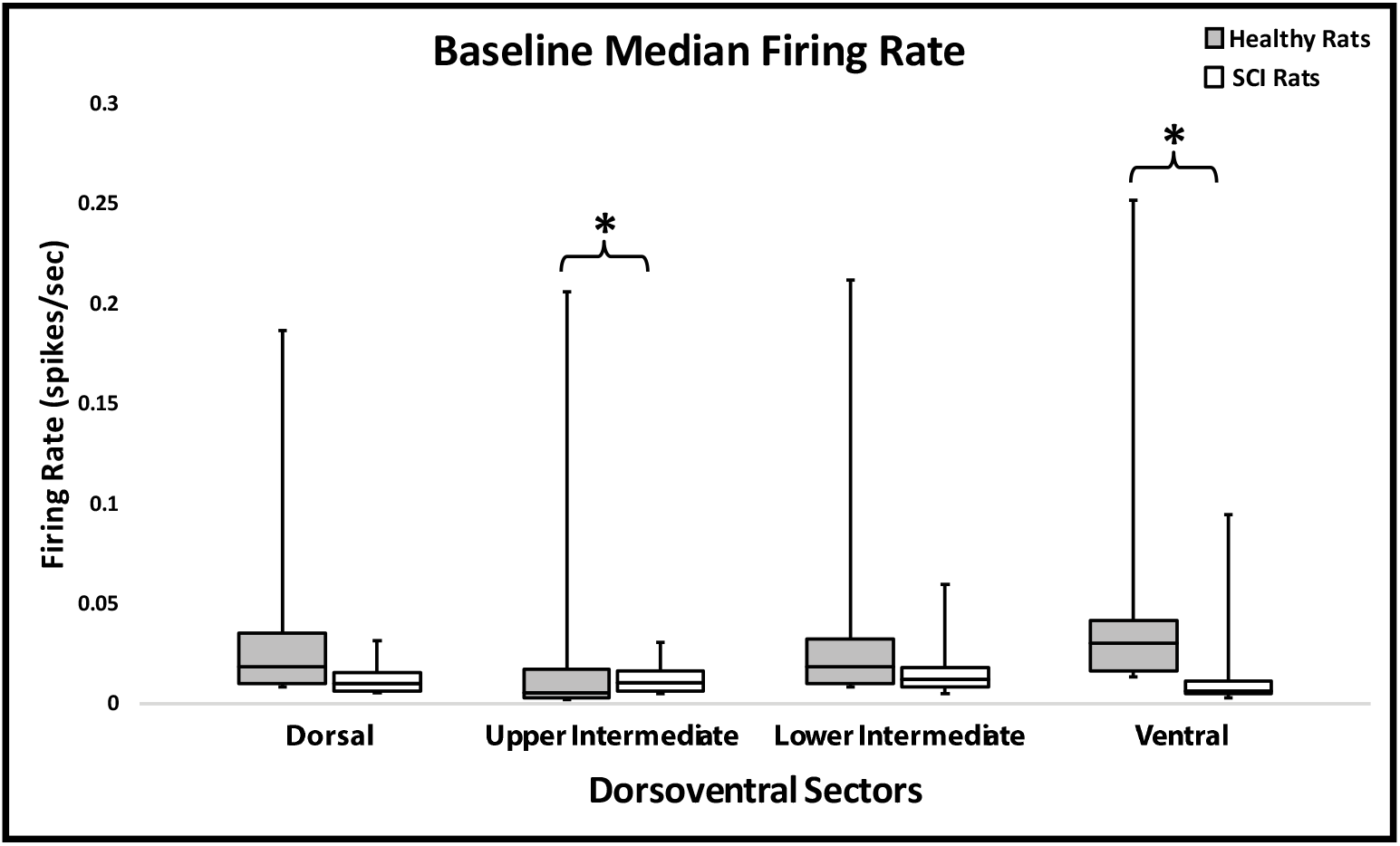
Baseline median firing rates of spontaneous spiking activity before onset ICMS (−20 ms to 0 ms spiking bins from post-stimulus spike histograms) in dorsoventral sectors based on depth below the surface of the spinal cord. The 16 recording channels were divided into 4 sectors based on depth below the surface of the spinal cord. **\***= Significant difference between healthy and SCI rats in the same dorsoventral sector (*p* < 0.05).

#### ICMS-evoked firing rate

The median firing rate of ICMS-evoked spikes was calculated for Healthy and SCI rats and is displayed in Figure 9. When comparing Healthy and SCI rats, the median firing rate of ICMS-evoked spikes at Short, L_on_, and L_p_ latencies generally decreased. The median firing rate of ICMS-evoked spikes at Short latencies was significantly lower in the Lower Intermediate (*p* = 0.0031) and Ventral (*p* = 0.0009) Sectors after SCI (Figure 9A). The median firing rate of ICMS-evoked spikes at L_on_ latencies was significantly lower in only the Upper Intermediate Sector (*p* = 0.0.0472) after SCI (Figure 9B). The median firing rate of ICMS-evoked spikes at L_p_ latencies was significantly lower in only the Ventral Sector (*p* = 0.0077) after SCI (Figure 9C).

**Figure 9.**
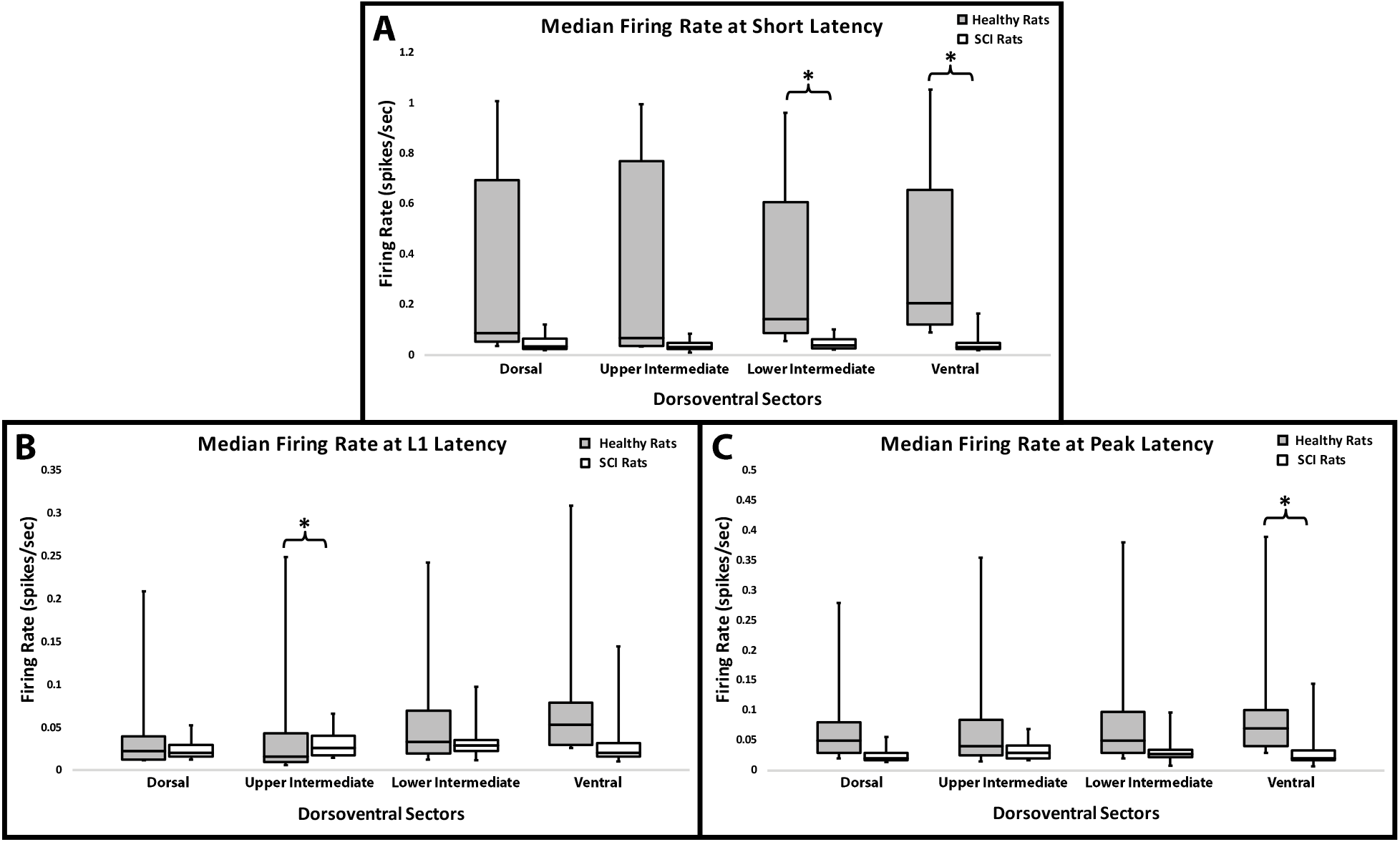
Median firing rates of ICMS-evoked spikes at specified latencies in dorsoventral sectors based on depth below the surface of the spinal cord. The 16 recording channels were divided into 4 sectors based on depth below the surface of the spinal cord. Median firing rates of ICMS-evoked spikes: at **A)** Short, **B)** L_ON_, and **C)** L_P_ latencies. **\***= Significant difference between healthy and SCI rats in the same dorsoventral sector (*p* < 0.05).

#### Spike amplitude

The median peak-to-peak amplitude of ICMS-evoked spikes recorded in the hindlimb spinal cord of Healthy and SCI rats are shown in Table 2. When comparing spike amplitudes between dorsoventral sectors in Healthy rats, the median ICMS-evoked spike amplitude was significantly higher in the Ventral Sector than in the Dorsal (*p* = 0.0043), Upper Intermediate (*p* = < 0.0001), and Lower Intermediate (*p* = 0.0013) Sectors. When comparing the median ICMS-evoked spike amplitudes of each dorsoventral sector between Healthy and SCI rats, the average ICMS-evoked spike amplitude was significantly lower in the Dorsal (*p* < 0.0001) and Ventral (*p* < 0.0001) Sectors.

**Table 2.** Median ICMS-evoked spike amplitude in healthy and SCI rats. Spike amplitude was calculated from peak-to-peak of the spike profile. The median spike amplitude was calculated for each dorsoventral sector. Bold and * = significantly lower (*p* < 0.05) when compared to healthy rats of the same dorsoventral sector.

| **Median ICMS-Evoked Spike Amplitude (mV)** |  |  |  |  |  |  |
| --- | --- | --- | --- | --- | --- | --- |
| **Dorsoventral Sector** | **Rat Group** | **10%** | **1<sup>st</sup> Quantile** | **Median** | **3<sup>rd</sup> Quantile** | **90%** |
| **Dorsal** | **Healthy** | 0.042 | 0.051 | 0.097 | 0.152 | 0.242 |
|  | **SCI** | 0.032 | 0.037 | **0.047\*** | 0.089 | 0.152 |
| **Upper Intermediate** | **Healthy** | 0.040 | 0.046 | 0.084 | 0.141 | 0.215 |
|  | **SCI** | 0.042 | 0.050 | 0.064 | 0.096 | 0.163 |
| **Lower Intermediate** | **Healthy** | 0.047 | 0.057 | 0.083 | 0.141 | 0.297 |
|  | **SCI** | 0.050 | 0.060 | 0.078 | 0.100 | 0.145 |
| **Ventral** | **Healthy** | 0.057 | 0.066 | 0.084 | 0.177 | 0.369 |
|  | **SCI** | 0.047 | 0.056 | **0.067\*** | 0.090 | 0.119 |

#### Spike latencies at different dorsolateral depths

The latencies of ICMS-evoked spikes recorded in the hindlimb spinal cord of Healthy and SCI rats are shown in Figure 10. When comparing the median ICMS-evoked spike latencies of each dorsoventral sector between Healthy and SCI rats, short latencies (Figure 10A) were significantly shorter in SCI rats in the Upper Intermediate (*p* = 0.0335) Sector. L_on_ latencies (Figure 10B) were significantly longer in SCI rats in the Upper (*p* = 0.0003) Sector. L_p_ latencies (Figure 10C) were significantly shorter in SCI rats in the Ventral Sector (*p* = 0.0179).

**Figure 10.**
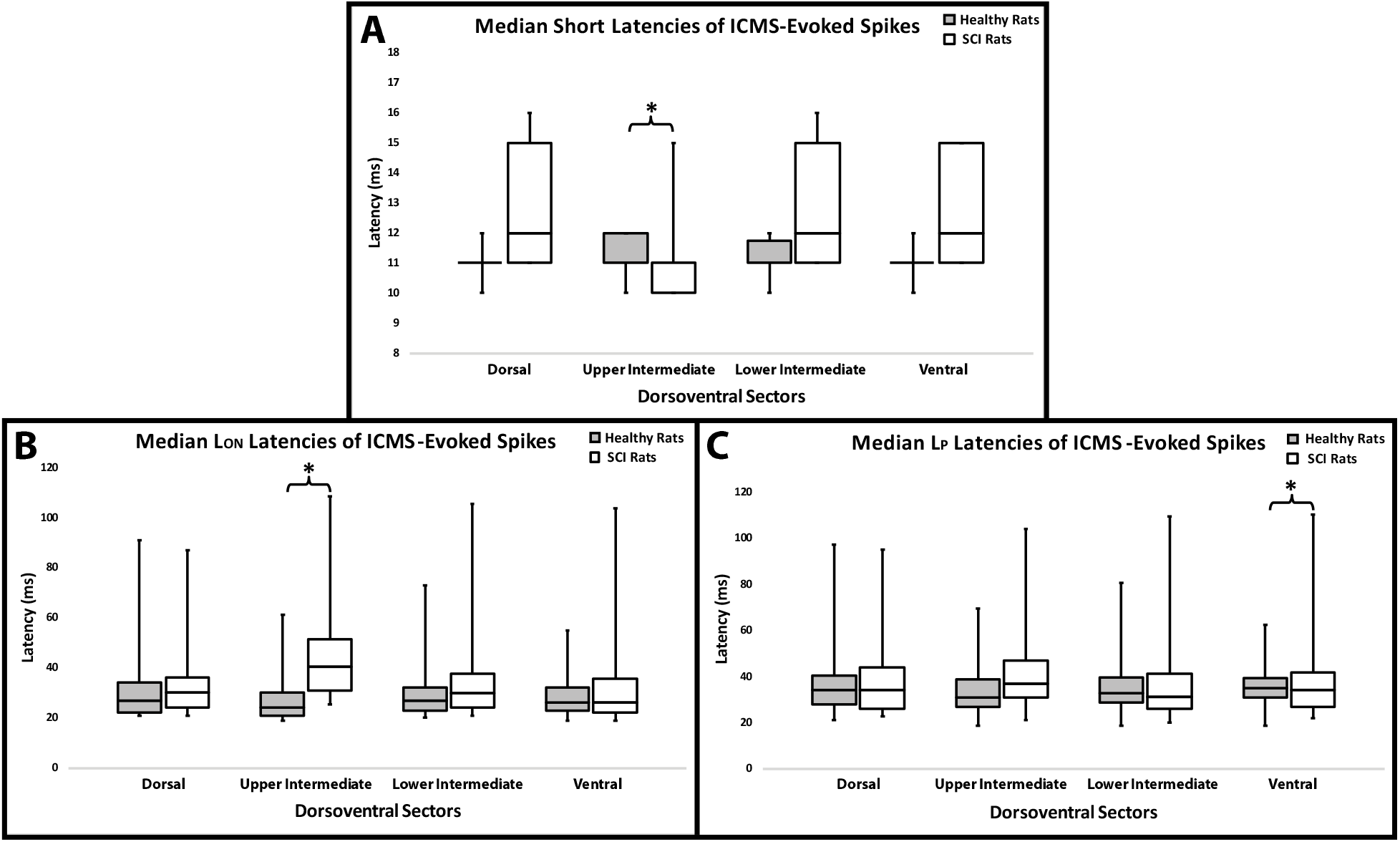
Median latencies of ICMS-evoked spikes by dorsoventral depth. Median latencies of ICMS-evoked spikes: at **A)** Short, **B)** L_on_, and **C)**, L_p_ latencies. **\***= Significant difference between healthy and SCI rats in the same dorsoventral sector (*p* < 0.05).

## Discussion

The goal of this study was to determine the effects of a contusive SCI on spinal motor neuron activity, corticospinal coupling, and conduction time in rats at various dorsoventral depths of the hindlimb spinal cord. In Healthy rats, ICMS-evoked activity in spinal cord neurons exhibited a distinct population of short-latency spikes reliably evoked at 10-12 ms at all dorsoventral depths with the exception of the Upper Intermediate Sector of the spinal cord. Four weeks after a contusion injury, severely damaging the CST in the dorsal columns, the short latency spikes were still present. Here we discuss the likely pathways underlying these short-latency responses and implications for understanding of how spared descending fibers participate in motor control after SCI.

### Functional Recovery after SCI in Rats

It has been shown in rats that synaptic coupling may be dysregulated after SCI, leading to hyperreflexia and spasticity (Yates et al., 2011). Immediately after SCI in humans, the spinal cord enters a state of spinal shock, a loss of sensation accompanied by motor paralysis, followed by a gradual recovery of reflexes (Ditunno et al., 2004). This is evident in this rat model of contusive SCI, where there is a loss of hindlimb motor function for roughly three days after injury, with reflexes and some voluntary use of hindlimbs returning over time. It has been reported that the duration and severity of spinal shock lasts up to two weeks in the rat (Bennett et al., 1999). Recovery of voluntary function plateaus at four weeks post-injury, but after a thoracic contusion deficits in control of hindlimb movements remain, as shown in this and a previous study (Krizsan-Agbas et al., 2014).

### ICMS-Evoked Neuronal Spikes in the Spinal Cord of Healthy Rats

Following ICMS in the HLA motor cortex, spikes were recorded reliably at all dorsoventral depths of the grey matter in the lower thoracic spinal cord. ICMS-driven spikes theoretically occur via one of three major routes: corticospinal tract (CST; either monosynaptic or disynaptic via spinal cord interneurons), cortico-rubro-spinal pathway, cortico-reticulo-spinal pathway, and cortico-vestibulo-spinal pathway. Based on antidromic activation studies, CST axons to the thoracic cord of rats have a maximum conduction velocity of ~19 m/s (average = 11 m/s) (Mediratta and Nicoll, 1983). Since the distance between the CST soma and the T13-L1 microelectrodes is ~20 cm, and assuming the *maximum* conduction velocity and a synaptic delay of 0.75 ms (Katz and Miledi, 1965), one would predict that the earliest spikes resulting from CST activation could occur at about 11 ms. However, based on the *average* conduction velocity (11 m/s), the CST-based spike distribution would peak at ~19 ms. Assuming a disynaptic connection (Alstermark et al., 2004), the earliest CST-based spikes would occur at ~12 ms and the peak at ~20 ms. Based on these assumptions, it is unlikely that the CST contributed substantially to the short-latency spike population in the present data. However, the long-latency spikes may be, at least in part, a result of disynaptic and polysynaptic CST connections and demonstrated by higher firing rates in intermediate and ventral laminae of the spinal cord grey matter. EMG latencies were observed beginning at 26-27 ms. Long-latency spikes recorded after the EMG response onset are likely intermixed with spike responses from Ia afferents subsequent to muscle contraction.

ICMS-evoked spikes also could have resulted from indirect activation via the cortico-rubrospinal, cortico-reticulospinal and cortico-vestibulospinal pathways. Rubrospinal tract (RbST) fibers have a similar conduction velocity to those of the CST (Al-Izki et al., 2008), so cortico-rubrospinal activity would at least partially overlap that of the corticospinal activity (with an additional synaptic delay) and could contribute to the long-latency response. Since the corticospinal and rubrospinal pathways are coupled with the evoked EMG activity, the CST and RbST may have a more direct influence on voluntary motor function, such as overground locomotion (DiGiovanna et al., 2016). But again, the longer latency responses are more difficult to interpret due to recurrent activation via Ia afferents.

Finally, while few studies have examined the conduction velocity of the reticulospinal tract (RtST) in rats, it appears that RtST conduction velocities are broadly distributed, but generally much faster than the CST or RbST in rats (Fox, 1970), and are similar to other species (Takakusaki et al., 2016). Conduction velocities of RtST neurons range from 16-80 m/s with a mean conduction velocity of 37 m/s (Fox, 1970). Thus, the earliest cortico-reticulospinal-based spikes could have occurred at T13-L1 at 4-5 ms. If spikes occurred at this time point, they could not be discriminated due to overlap with the stimulus artifact. Based on the mean conduction velocity, spikes would occur at about 8 ms. But due to the wide range of conduction velocities expected in the RtST, it is likely that the short-latency responses at 10-12 ms were primarily due to cortico-reticulospinal fiber activation. This pathway is assumed to have disynaptic or polysynaptic connections either directly onto long-propriospinal neurons or commissural interneurons located in the grey matter (Mitchell et al., 2016). In the present study, firing rates were similar throughout the dorsoventral layers. To our knowledge, studies have not been conducted measuring the conduction velocity of the vestibulospinal, uncrossed ventral corticospinal, and crossed dorsolateral corticospinal tracts in the rat, so direct comparisons cannot be made.

### ICMS-Evoked Neuronal Spikes in the Spinal Cord of SCI Rats

A moderate contusion applied to the dorsal midline of the thoracic spinal cord (modeling a clinical injury in humans) destroys the central core of the spinal cord at the lesion epicenter, ablating the spinal grey matter and sparing a limited amount of circumferential descending motor pathways (Basso et al., 1996; Lemon, 2008; Fink and Cafferty, 2016). Specifically, the dorsomedial funiculi are severely damaged, while the dorsolateral, ventral, and ventromedial funiculi remain largely intact.

The mean baseline activity before ICMS onset may be indicative of the number of viable fibers terminating in each dorsoventral laminae. After SCI, the median baseline spiking activity was decreased in the upper intermediate and ventral laminae (Figure 8). This may suggest that the number of viable fibers terminating in these laminae decreased after SCI while the fibers terminating in the dorsal and lower intermediate laminae remained relatively undamaged. However, without a tract-tracing analysis, it is difficult to correlate this activity to any specific descending or ascending pathway. Further tract-tracing analysis is needed to verify this assumption. In addition, baseline activity could be related to the level of anesthesia during the recording procedure. However, the average amount of ketamine administered was not significantly different between groups, suggesting that the level of anesthesia was not a contributing factor in the difference in baseline spiking activity between groups.

Most notably, short-latency spikes were still present despite severe damage to the CST. This is most likely due to the maintenance of the cortico-reticulospinal tract. Although the firing rate was diminished (Figure 9A), the probability of activation (POA) for short-latency spikes was increased in SCI rats in all dorsoventral laminae, roughly doubling in dorsal, upper intermediate and ventral laminae, and nearly tripling in the lower intermediate laminae (Table 1). This suggests that in healthy rats the dorsomedial CST exerts an inhibitory influence on spinal cord neurons, and that after SCI, ICMS-evoked spike activity via other descending pathways is released from inhibition. This remaining short-latency spike activity provides a putative substrate for the effects of spared descending spinal fibers to influence spinal cord function after injury.

Additionally, after injury, the median conduction time of the short-latency spikes was the same in the dorsal, lower intermediate, and ventral laminae but slightly shorter in the upper intermediate laminae. This is strong evidence that the spared cortico-reticulospinal fibers terminating in the dorsal lower intermediate, and ventral layers remained unaltered after injury (Figure 10). Furthermore, forelimb-hindlimb and left-right coordination is controlled via long descending propriospinal pathways (Frigon, 2017), which receive a larger number of terminals from reticulospinal neurons as compared to corticospinal neurons, suggesting that left-right activity is strongly controlled by the RtST (Mitchell et al., 2016). At 4 weeks post-SCI, rats recovered coordination as noted by their BBB scores, suggesting that RtST remained intact.

After SCI, the long-latency (L_on_ and L_p_) spikes were still present, but firing rate was significantly increased in intermediate (L_p_; Figure 9B) and significantly diminished in ventral (L_on_; Figure 9C) laminae. The L_on_ latency was longer in the intermediate laminae (Figure 10B) while the L_p_ latency was shorter in the ventral laminae (Figure 10C). But, the POA was increased in the lower intermediate (L1) and ventral laminae (L1 and peak) and decreased in the upper intermediate laminae (peak) (Table 1). The unaltered spiking activity and POA in the dorsal laminae suggests that the CST and RbST have a relatively minor influence on the dorsal laminae, at least in the lumbar spinal cord, and that any influence of the RtST on the dorsal laminae remains unchanged.

In rats, the injured CST has been shown to sprout at cervical locations in response to a bilateral transection of the thoracic spinal cord, adding novel connections onto long propriospinal neurons that may transmit signals to denervated neurons below the injury (Fouad et al., 2001; Bareyre et al., 2004). The delayed conduction times and increased POA could be evidence of corticospinal fibers sprouting onto long propriospinal neurons that bridge the lesion, which in turn could have a greater influence on spinal cord neurons below the lesion in intermediate and ventral laminae (Bareyre et al., 2004). However, the conduction times of the peak long-latency response was significantly shorter in the ventral laminae. It is possible that the disruption of CST fibers resulted in disinhibited reticulospinal influence on spinal cord neurons in the ventral laminae. Furthermore, plasticity of other pathways could play a role in these changes, such as the dorsolateral (Hilton et al., 2016) and ipsilateral ventral (Bareyre et al., 2004) CST.

#### Functional Significance for Recovery after SCI

Over time, rats with moderate thoracic spinal cord contusions naturally recover some hindlimb motor function but deficits are still present, as observed in the lower BBB scores 4 weeks post-SCI in the present study. With remaining deficits, further rehabilitation may be beneficial for a greater recovery of motor function. The results of this study indicate an intact reticulospinal pathway after moderate contusion could be a target for rehabilitation. Targeted therapy of an intact reticulospinal tract in rats with severe spinal cord contusion has been shown to result in the reorganization of the cortico-reticulo-spinal circuit and improvement of motor function (Asboth et al., 2018). Stimulation therapies targeting specific descending motor pathways using time-dependent stimulation techniques have shown substantial recovery of motor function after injury in humans (Bunday et al., 2018; Sangari et al., 2019). Understanding the conduction latencies of intact pathways after injury may aid in the development of non-invasive stimulation therapies that can be translated to the clinic. This understanding of conduction latencies, corticospinal coupling, and intact descending pathways after injury is important for future neuromodulatory therapies which could lead to robust recovery after spinal cord contusion.

## Conclusion

The effects of a contusive SCI on spinal motor neuron activity, corticospinal coupling, and conduction time in rats were presented and described. ICMS-evoked spiking activity decreased after SCI; however, the presence of ICMS-evoked spikes at short latencies (10-12 ms) after SCI indicates that cortico-reticulospinal fibers, with their faster conduction velocities, remain intact after a moderate contusive injury, and CST damage may release their effects from inhibition. These data inform the further development of latency specific stimulation therapies and neuromodulation techniques in the lumbar spinal cord for recovery after SCI.

## Acknowledgments

This work was supported by the Paralyzed Veterans of America Research Foundation #3068, The Ronald D. Deffenbaugh Family Foundation, NIH/NINDS R01 NS030853, T32 Neurological and Rehabilitation Sciences Training Program, and NIH/NINDS F31 NS105442.

